## Supplementary Information for "Machine-learning-based motion tracking revealed the inverse correlation between adhesivity and surface motility of the leptospirosis spirochete"

Table S1. Bacterial strains used in this study.

Table S2. Summary of experimental data.

Figure S1. Swimming and crawling speeds of the mutant strains on NRK.

Figure S2. Adherence and motility of the mutant strains on MDCK.

Figure S3. Transposon insertion mutants.

Figure S4. Effect of GMM parameters on the detection of paused bacteria.

Figure S5. Dirichlet distribution.

Movie S1. Non-processed video of leptospires over kidney cells (Figure 2a, Step1).

Movie S2. Background subtraction (Figure 2a, Step2).

Movie S3. Morphological transformation (Figure 2a, Step3).

Movie S4. Sorting cells from non-cells (Figure 2a, Step4).

Movie S5. The result of tracking crawling bacteria (Figure 2a, Step5).

**Table S1. Bacterial strains used in this study.**

| Bacterial strain | | | | | Abbr. | Isolated from | Ref. |
|---|---|---|---|---|---|---|---|
| *L. interrogans* | | | | | | | |
| serogroup | Hebdomadis | strain | D-OW16-2K | | Hebd-16 | Dog | This study |
| serogroup | Hebdomadis | strain | D-SO21-6K | | Hebd-21 | Dog | This study |
| serogroup | Australis | strain | D-FO15-8K | | Australis | Dog | This study |
| serogroup | Icterohaemorrhagiae | strain | RN-TK-AS1707-6 | | Ictero | Rat | This study |
| serogroup | Canicola | strain | D-NA10-E1-0K | | Canicola | Dog | This study |
| serogroup | Autumnalis | strain | D-SA11-5K | | Autum | Dog | [1] |
| serovar | Manilae | strain | UP-MMC-NIID | | Manilae | Human | [2] |
| serovar | Manilae | strain | UP-MMC-NIID | *ligA::Tn* | *ΔligA* | - | This study |
| serovar | Manilae | strain | UP-MMC-NIID | *lenA::Tn* | *ΔlenA* | - | This study |

**Table S2. Sammary of adhesion and motility assays.**

| Host Kidney cells | Bacteria *Leptospira* | Adhesion to kidney cells Bacteria/mm$^2$ (fields) | Swimming speed $v_{SW}$ (μm/s) (bacteria) | Crawling speed $v_{CR}$ (μm/s) (bacteria) | Crawling ability $v_{CR}$ / $v_{SW}$ |
|---|---|---|---|---|---|
| NRK (Rat) | Hebd-16 | 2899 ± 198 (n = 16) | 6.3 ± 0.3 (n = 93) | 3.7 ± 0.1 (n = 124) | 0.6 ± 0.1 |
| | Hebd-21 | 2587 ± 221 (n = 18) | 12.8 ± 0.3 (n = 200) | 7.3 ± 0.5 (n = 100) | 0.6 ± 0.1 |
| | Australis | 1575 ± 157 (n = 14) | 5.5 ± 0.2 (n = 131) | 5.0 ± 0.2 (n = 120) | 0.9 ± 0.1 |
| | Ictero | 2647 ± 177 (n = 12) | 9.7 ± 0.0 (n = 329) | 6.4 ± 0.2 (n = 432) | 0.7 ± 0.0 |
| | Canicola | 3268 ± 249 (n = 12) | 10.1 ± 0.4 (n = 112) | 3.8 ± 0.2 (n = 79) | 0.4 ± 0.1 |
| | Autum | 1108 ± 66 (n = 16) | 3.7 ± 0.1 (n = 220) | 6.6 ± 0.3 (n = 119) | 1.8 ± 0.1 |
| | Manilae | 3582 ± 367 (n = 16) | 6.9 ± 0.2 (n = 213) | 4.7 ± 0.2 (n = 190) | 0.7 ± 0.0 |
| | *ΔlenA* | 1546 ± 117 (n = 20) | 5.0 ± 0.2 (n = 161) | 5.4 ± 0.2 (n = 116) | 1.1 ± 0.1 |
| | *ΔligA* | 1533 ± 111 (n = 15) | 6.2 ± 0.2 (n = 114) | 4.9 ± 0.2 (n = 108) | 0.8 ± 0.1 |
| MDCK (Dog) | Hebd-16 | 680 ± 133 (n = 14) | 4.3 ± 0.2 (n = 113) | 5.7 ± 0.3 (n = 117) | 1.3 ± 0.1 |
| | Hebd-21 | 1489 ± 211 (n = 15) | 11.7 ± 0.3 (n = 192) | 10.2 ± 0.3 (n = 170) | 0.9 ± 0.0 |
| | Australis | 1760 ± 245 (n = 12) | 6.2 ± 0.2 (n = 250) | 5.0 ± 0.3 (n = 116) | 0.8 ± 0.1 |
| | Ictero | 1621 ± 145 (n = 15) | 10.2 ± 0.3 (n = 224) | 9.6 ± 0.2 (n = 256) | 0.9 ± 0.0 |
| | Canicola | 1753 ± 252 (n = 15) | 7.5 ± 0.3 (n = 103) | 8.1 ± 0.5 (n = 110) | 1.1 ± 0.1 |
| | Autum | 528 ± 102 (n = 20) | 5.0 ± 0.3 (n = 116) | 7.2 ± 0.2 (n = 190) | 1.4 ± 0.1 |
| | Manilae | 1005 ± 117 (n = 18) | 6.3 ± 0.2 (n = 188) | 6.8 ± 0.2 (n = 194) | 1.1 ± 0.0 |
| | *ΔlenA* | 1032 ± 173 (n = 18) | 3.9 ± 0.1 (n = 177) | 5.7 ± 0.2 (n = 123) | 1.4 ± 0.0 |
| | *ΔligA* | N.D. | 7.5 ± 0.4 (n = 72) | N.D. | N.D. |

See Figure S2 for N.D. of Δ*ligA*.

### Effect of the loss of functional outer membrane protein genes on adhesin and motility

We measured free-swimming speeds of the wild-type (WT) of *L. interrogans* serovar Manilae and the transposon-insertion mutants ($\Delta lenA$ and $\Delta ligA$) and found that the disruption of the genes significantly affected the swimming speed (Figure S1). Therefore, we used the averaged crawling speeds normalized by the averaged swimming speeds to evaluate the crawling ability of *Leptospira* strains. The value represented by $v_{\mathrm{CR}}/v_{\mathrm{SW}}$ in the main text is the ratio of the speed given by the cell rotation when attached to the host cells to the speed when detached. The errors of $v_{\mathrm{CR}}/v_{\mathrm{SW}}$ were calculated from relative errors of crawling speeds and swimmng speeds using a general equation of error propagation: the relative error of $v_{\mathrm{CR}}/v_{\mathrm{SW}}$ is determined by quadratic sum of relative errors of $v_{\mathrm{CR}}$ and $v_{\mathrm{SW}}$. As shown in Figure 4a, swimming speeds were different among strains. We thus used $v_{\mathrm{CR}}/v_{\mathrm{SW}}$ to compare the crawling ability among strains in Figure S2, and Figures 5 and 6 in the main text.

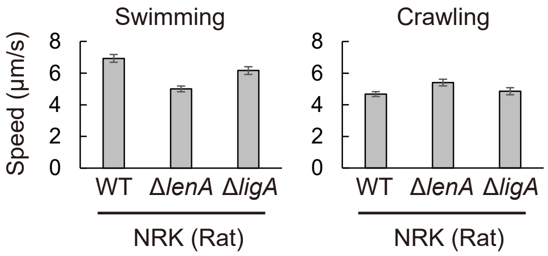


Figure S 1. Swimming and crawling speeds of WT and mutant strains of *L. interrogans* serovar Manilae on NRK. Average values and standard errors are shown. The number of bacteria measured is shown in Table S2.

We also measured adherence and motility of the WT and mutant strains of *L. interrogans* serovar Manilae over the dog kidney cells MDCK. $\Delta lenA$ retained adhesivity, and its crawling ability was higher than WT (Figure S2). In contrast, very few $\Delta lenA$ cells attached to MDCK, and thus no motiity data were obtained (indicated as N.D. in Figure S2). These results suggest that LenA would be involved in attachment and crawling on MDCK, but LigA likely plays more critical role.

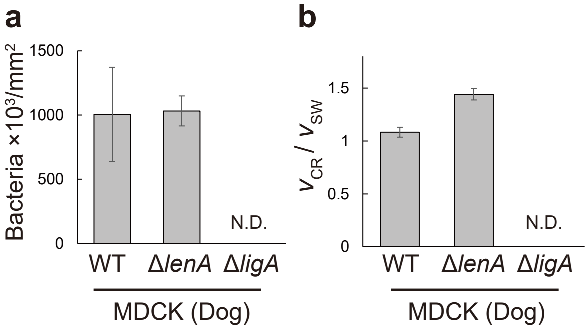


Figure S 2. Adhesion (a) and crawling ability (b) of WT and mutant strains of *L. interrogans* serovar Manilae on MDCK. No data (N.D.) for $\Delta ligA$ strain.

### Transposon insertion mutants

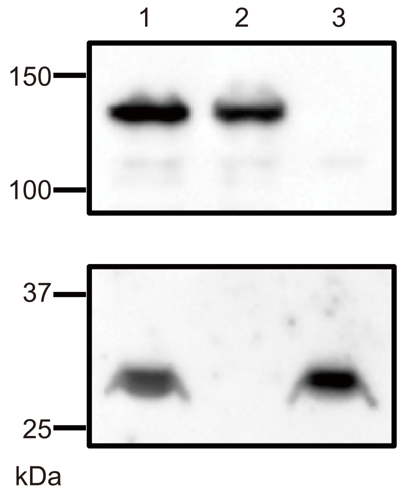


Figure S 3. Immunoblot analysis of whole cell lysates from *L. interrogans* wild-type and transposon insertion mutants. *L. interrogans* whole cell lysates ($1.5 \times 10^8$ cells) were subjected to 5–20% SDS-PAGE and Western blotting with anti-LigA (upper panel) or anti-LenA (lower panel). Lanes 1, wild-type; 2, *lenA*::*Tn*; 3, *ligA*::*Tn*.

### The theoretical limit of bacteria tracking

As explained in the main text, pixel-value distributions are modeled by the Gaussian mixture model (GMM):

$$\hat{p}(x|H_T, \mathrm{BG}+\mathrm{FG}) = \sum_{\mathrm{m}=1}^{\mathrm{M}} \hat{\pi}_{\mathrm{m}} \mathcal{N}(x; \hat{\mu}_{\mathrm{m}}, \hat{\sigma}^2_{\mathrm{m}}). \tag{1}$$

This model provides $M$ Gaussian distributions, individually weighted by $\hat{\pi}_m$ ($\sum_{m=1}^{M} \hat{\pi}_m = 1$). In the background subtraction, the fractions of foreground (FG) and background (BG) should be determined arbitrarily. If the fraction of FG is defined as $c_{\mathrm{fg}}$, that of BG is $1 - c_{\mathrm{fg}}$. The present study assumes $1 - c_{\mathrm{fg}} \ll c_{\mathrm{fg}}$. Therefore, when $\hat{\pi}_m$ is added from the largest one in descending order until it exceeds $1 - c_{\mathrm{fg}}$, the result indicates the total fraction of BG:

$$B = \arg\min_{\boldsymbol{bg}} \left( \sum_{m=1}^{bg} \hat{\pi}_m > (1 - c_{\mathrm{fg}}) \right), \tag{2}$$

where $B$ represents the number of clusters comprising BG distribution. Using $B$, BG distribution can be modeled by:

$$\hat{p}(x|H_T, \mathrm{BG}) \sim \sum_{\mathrm{m}=1}^{\mathrm{B}} \hat{\pi}_{\mathrm{m}} \mathcal{N}(x; \hat{\mu}_{\mathrm{m}}, \hat{\sigma}^2_{\mathrm{m}}). \tag{3}$$

Consider the case of FG cluster generation when a new pixel value obtained upon object intruding is determined not to belong to any existing clusters. If the object comes to a halt, $\hat{\pi}_m$ is increased with the dwell time according to the following update equation (Eq. 2 in the main text),

$$\hat{\pi}_m \leftarrow \hat{\pi}_m + \alpha(o_m^{(t)} - \hat{\pi}_m) - \alpha c_T, \tag{4}$$

where $c_T = -c_m/T$ (see "Dirichlet distribution" in this Supplementary Information). If $\hat{\pi}_m$ is increased for $n$ frames because of changeless pixel value (i.e., the object stays the same place for $n$ frames), Eq. (4) indicates that $\hat{\pi}_m$ reaches $1 - (1-\alpha)^n$ (assuming $c_T = 0$). When the weight exceeds FG fraction $c_{\mathrm{fg}}$, $1 - (1-\alpha)^n > c_{\mathrm{fg}}$, the FG object is misrecognized. Namely, $\alpha$ and $c_{\mathrm{fg}}$ are key parameters determining the theoretical limit $n$, discriminating moving objects from BG. According to the inequality above, $n$ is determined by:

$$n > \frac{\log(1 - c_{\mathrm{fg}})}{\log(1-\alpha)}. \tag{5}$$

For example, when $c_{\mathrm{fg}} = 0.1$ and $\alpha = 0.001$, $n$ ∼100 frames. This indicates that, even if the object should be FG, it is recognized as BG by halting exceeding 100 frames. Figure S4 shows the effect of $n$ on the FG (bacteria) detection, demonstrating that the increased $n$ enhances noise due to the misrecognition of FG, which prevents motion tracking. Such a problem can be solved by increasing $c_{\mathrm{fg}}$ or decreasing $\alpha$. However, given that $\alpha$ is a modulator decaying the contribution of past data, a decrement of $\alpha$ affects the rate of parameter update adaptation against scene changes. Moreover, the increased $c_{\mathrm{fg}}$ could promote the misrecognition of FG as BG. Hence, appropriate values of $c_{\mathrm{fg}}$ and $\alpha$ should be verified based on the locomotion property of bacteria. The motion speed is a significant factor to warrant adequate parameter setting. The previous study defined the measured values less than 1 µm/s as non-motile data [3]. Consider a bacterium moving at 1 µm/s. If the cell length is 8 µm (an average length), the bacterium takes 8 s to pass through a pixel. In this study, we recorded bacteria at 0.2 µm/pixel, which is negligibly small against movement distance. Given that the recording frame rate was 30 fps, $n = 240$ is required to detect motion faster than 1 µm/s. Based on these considerations, we set $c_{\mathrm{fg}} = 0.22$ and $\alpha = 0.001$ ($n$ ∼250 frames). Note that bacteria stopping for longer than 8 s will be removed from tracking targets. Hence, our methods can recognize bacteria moving at the speed of >1 µm/s and not stopping for >8 s.

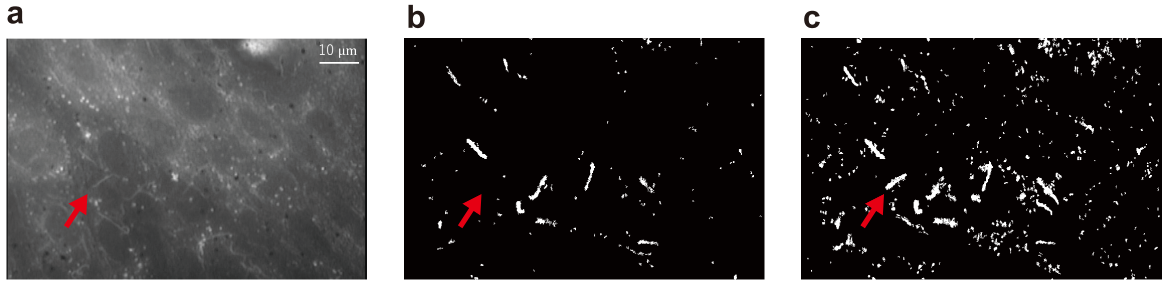


Figure S 4. Effect of GMM parameters on detection of paused bacteria. (a) Raw image. The bacteria indicated by the red arrow has been stopped for more than 100 seconds. The bacterium was not detected by $c_{\mathrm{fg}} = 0.1$ and $\alpha = 0.01$ (n∼10) (b), whereas it was detected by $c_{\mathrm{fg}} = 0.1$ and $\alpha = 0.001$ (n∼105) (c).

### Dirichlet distribution

The Dirichlet distribution is the conjugate prior of the multinominal distribution:

$$P(\vec{x};\vec{\alpha}) = \frac{1}{Z}\prod_{i=1}^{K} x_i^{\alpha_i - 1}, \tag{6}$$

where $Z$ is beta function, $\vec{x} = (x_1, x_2, \sim x_{K-1}, x_K)$ is the stochastic variable ($x_i \geq 0$ and $\sum_{i=1}^{K} x_i = 1$), $\vec{\alpha} = (\alpha_1, \alpha_2, \sim \alpha_{K-1}, \alpha_K)$ is the hyper parameter ($\alpha > 0$). When $K$ events independently occur $\alpha_i - 1$ times for each independently, the distribution gives the occurrence probability of $\vec{x}$.

Consider a three-sided dice numbered from 1 to 3. After rolling the dice many times, the probability at which the dice shows each pip gets close to the same probability: $\vec{x} = (1/3, 1/3, 1/3))$. This means that rolling the dice 30 times results in the occurrence of 10 times for each pip. However, repeating the trial will not show $\vec{x} = (1/3, 1/3, 1/3))$ every time. Namely, though $\vec{x} = (1/3, 1/3, 1/3)$ is the most likely outcome, an infinit number of patterns exist in a range satisfying $\sum_{i=1}^{K} x_i = 1$. This example, assuming that $\alpha$ is a uniform distribution, can be modeled by the Dirichlet distribution with $\alpha > 1$ (Figure S5a). The distribution shows a peak at the center of the triangle, that is $\vec{x} = (1/3, 1/3, 1/3)$. In this study, we assumed that the background (cultured cells) cluster accounts for a major portion of the distribution, and the foreground (bacteria passing through the pixel) is minor. Such strongly biased Dirichlet distribution is given by $0 < \alpha < 1$, biasing $x_i$ to 0 or 1 because of $x_1^{\alpha_1 - 1} x_2^{\alpha_2 - 1} x_3^{\alpha_3 - 1} \to 0$ (Figure S5b). $\alpha_i - 1$ corresponds to $\hat{\pi}_m^{c_m}$ of Eq. (7) in the main text. Therefore, we set $-1 < c_m < 0$.

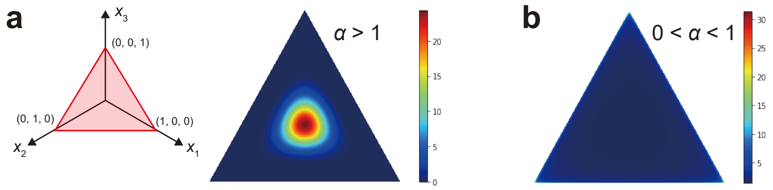


Figure S 5. Dirichlet distributions. (a) The stochastic variables $\vec{x}$ exist within the red triangle because of $\sum_{i=1}^{K} x_i = 1$ (left). The distribution simulated with $\alpha = 10$ (right). (b) $\alpha = 0.9$.
